# A Joint Subspace Mapping Between Structural and Functional Brain Connectomes

**DOI:** 10.1101/2022.04.12.488055

**Authors:** Sanjay Ghosh, Ashish Raj, Srikantan S. Nagarajan

## Abstract

Understanding the connection between the brain’s structural connectivity and its functional connectivity is of immense interest in computational neuroscience. Although some studies have suggested that whole brain functional connectivity is shaped by the underlying structure, the rule by which anatomy constraints brain dynamics remains an open question. In this work, we introduce a computational framework that identifies a joint subspace of eigenmodes for both functional and structural connectomes. We found that a small number of those eigenmodes are sufficient to reconstruct functional connectivity from the structural connectome, thus serving as low-dimensional basis function set. We then develop an algorithm that can estimate the functional eigen spectrum in this joint space from the structural eigen spectrum. By concurrently estimating the joint eigenmodes and the functional eigen spectrum, we can reconstruct a given subject’s functional connectivity from their structural connectome. We perform elaborate experiments and demonstrate that the proposed algorithm for estimating functional connectivity from the structural connectome using joint space eigenmodes gives competitive performance as compared to the existing benchmark methods with better interpretability.

## 1. Introduction

The connection between the dynamics of neural processes and the anatomical substrate of the brain is a central question in neuroscience research. Understanding this interplay is essential for understanding how behaviour emerges from the underlying anatomy [1, 2]. To this extent, a common way to represent the whole brain is using a network or graph [3], where nodes represent cortical and sub–cortical gray matter volumes and edges stand for the strength of structural or functional connectivity. Structural connectivity is typically extracted from tractography algorithms applied to diffusion magnetic resonance imaging (MRI) or diffusion tensor imaging (DTI) data [4]. Functional connectivity usually refers to pairwise correlation between activation signals in various brain regions is measured by various functional brain imaging modalities - functional MRI (fMRI) [5], electroencephalography (EEG) [6], magnetoencephalography (MEG) [7] etc. Although several studies [8, 9, 10, 11, 12, 13, 14, 15, 16] have suggested that whole brain functional connectivity is shaped by the underlying structure, the rule by which anatomy constraints brain dynamics remains an open question and an interesting research challenge [10]. We note that a deeper analysis could explain how signals propagate within a structured graph between different regions, and interfere or interact with each other to induce a global pattern of temporal correlations. Conversely, a fuller understanding of this relationship will allow us to explore what portion of functional activity that are *not* explainable by structural connectome alone; potentially reflecting more complex transynaptic processing across brain networks [17, 18, 19].

Several studies have aimed towards understanding of the mapping between structural and functional connectivity [20, 17, 16, 21, 22, 19, 23, 24, 25]. A broad class of these studies are built on complex generative models of functional activity [10, 20, 26, 27, 28, 29, 25] typically that generate simulated functional time–series from which the functional connectome is estimated. In contrast, some direct approaches [21, 19, 30] rely on the eigen decomposition of the structural connectivity matrix or its Laplacian and finding a link to eigendecomposition of the functional connectivity matrix [31, 21, 32, 19, 30, 33, 34, 35]. Authors in [19] predict the functional connectome from the eigenmodes of structural Laplacian. In this paper, we use the term ”eigenmode” to refer eigenvectors and ”eigen spectrum” to refer eigenvalues. Similarly, authors in posed the structure-function mapping as a *L*_2_ minimization problem. However, similar to [19], the feasible eigenmodes were restricted to the individual eigenmodes of structural connectome (roughly equivalent to Laplacian eigenmodes). Despite the success of this approach, it can be limiting to enforce the functional eigenmodes to belong to the space of the structural Laplacian eigenmodes. Recently Becker et al. [30] introduced an idea of using rotational operator between the connectomes. Recently, authors in [36] introduced a unified framework by which most existing mappings based on eigenmodes can be expressed [See Sec. 2.4 in [36]]. In particular, the predicted functional connectome by the spectral mapping of [30] was shown to have a structured eigen polynomial of degree N, the number of ROIs. Moreover, to achieve improved predictions, the authors in [36] proposed to add a constant symmetric matrix term within the expression of predictive model. This structure-function mapping was posed was Riemannian distance [37] minimization problem in [38].

In this paper, we introduce a new approach to integrate both structural Laplacian and functional connectivity by a common vector space. The underlying neuroscientific assumption is that functional correlations arise due to signal transmission on the structural network, hence the two should share a common set of “modes” or eigenvectors. However, a one-way mapping from structure to function, may not fully capture all underlying biological processes. The structural network is biased towards monosynaptic connections and also may not be considered a gold standard in view of our ability to accurately measure all connections (especially lateral cortical connections). Therefore, there remains a need for a joint mapping approach that does not privilege one or the other connectome. We build upon the seminal work in [17, 18, 19] that investigated the use of brain connectivity harmonics as a basis to represent spatial patterns of cortical networks. By using the orthogonality of connectome harmonics, it is shown that a linear combination of these eigenmodes can be used to recreate any spatial pattern of neural activity.

This joint estimation framework allows us to express functional connectivity of a particular subject (brain sample) as a subspace of the joint eigenmodes of both structure and function. We demonstrate that just a small fraction of the proposed joint eigenmodes are sufficient to span the functional connectivity. We further extend the core idea to develop a predictive model that is able to accurately predict functional connectivity from structural Laplacian. The bottleneck for this prediction model is to find a mapping from the projection of structural Laplacian and functional connectome to the joint space. We circumvent this by using a nonlinear and data-driven mapping technique. We refer our proposed predictive method as *joint eigen spectrum mapping* (JESM). This finally gives us an efficient method for structure-function mapping of human brain connectivity. In summary, the present work emphasizes on a deeper theoretical foundation towards analyzing brain structure and function. Two main contributions in this paper are:

1. We demonstrate how to jointly diagonalize both structural and functional connectivities using a common set of eigenmodes. To the best of our understanding, this is an interesting finding in context human brain connectivity analysis. This exploration could serve as a building block for more future research on the impact of brain diseases on brain structural and functional connectivity.
2. We propose an optimization based method for structure-function mapping using joint eigenmodes and the relationship between joint eigen spectra of both the connectomes.

This paper is outlined as follows. In Section 2, we introduce our framework governing the relationship between structural Laplacian and functional connectivity. The details on structure-function predictive model are presented in Section 3. In Section 4, we perform thorough experimental analysis on a large dataset of healthy subjects and also compare our approach with existing state-of-the-art methods. We also experiment on data of schizophrenic patients to demonstrate that the proposed method generalizes beyond healthy subjects. Finally, we conclude in Section 5 with a summary of our work.

## 2. Theory and Methods

Suppose ***S*** ∈ ***ℝ***^*n×n*^ and ***F*** ∈ ***ℝ***^*n×n*^ are the structural and functional connectivity matrices of an arbitrary subject. Here *n* corresponds to the number of regions-of-interest (ROI) in a brain atlas. The entry (*i, j*) of these matrices represents the strength of the connectivity between regions *i* and *j* evaluated either structurally or functionally. In Table 1 we describe the parameters and variables used in this work. We consider the problem of finding a joint or common vector space between ***S*** and ***F***. We perform a unique decomposition of both connectomes sharing the same algebraic structure. This is referred as *simultaneous diagonalization*, which is a well-studied area in signal processing [40, 41, 42]. Our goal is to find a matrix *A* = [***a***_1_|***a***_2_| … |***a***_*n*_] and ***a***_*i*_ ∈ *ℝ*^*n×*1^, ∀*i* = {1, 2, …, *n*} that can express both structural and functional connectivity by its orthonormal column vectors ***a***_*i*_. Without loss of generality, we instead work with structural Laplacian of the structural connectivity as in [19]. Let ***D*** be the (diagonal) degree matrix of structural connectivity ***S***. Then, the (normalized) Laplacian of the structural connectivity is given by:

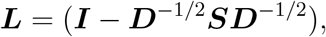

where ***I*** ∈ *ℝ*^*n×n*^ is an identity matrix. Finally, the revised mathematical problem is to find a matrix 𝒜 = [***a***_1_|***a***_2_| … ***a***_*n*_] such that:

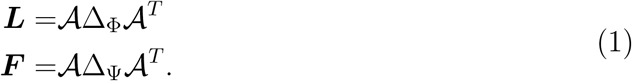

**Table 1:** Summary of the variables and definitions used in this text.

| Symbol | Description |
| --- | --- |
| $n$ | Number of region-of-interest (ROI). |
| $\mathbf{S}$ | Structural connectome of size $(n \times n)$. |
| $\mathbf{L}$ | Structural Laplacian. |
| $\mathbf{F}$ | Functional connectome. |
| $\Lambda$ | Eigenvalues (/eigen spectra) of structural Laplacian ($\mathbf{L}$). |
| $U$ | Eigenvectors of structural Laplacian. |
| $\Gamma$ | Eigenvalues of functional connectome. |
| $V$ | Eigenvectors (/eigenmodes) of functional connectome. |
| $\Phi$ | Joint eigenvalues of structural Laplacian ($\mathbf{L}$). |
| $\Psi$ | Joint eigenvalues of functional connectome ($\mathbf{F}$). |
| $\mathcal{A}$ | Joint eigenmodes of $\mathbf{L}$ and $\mathbf{F}$. |
| $m$ | Number of subjects. |
| $\mathbf{L}_j$ | Laplacian of subject $j$. |
| $\mathbf{F}_j$ | Functional connectome of subject $j$. |
| $\Phi_j$ | Joint eigenvalues of structural Laplacian of subject $j$. |
| $\Psi_j$ | Joint eigenvalues of functional connectome of subject $j$. |
| $\Delta_\Lambda$ | Diagonal matrix with diagonal entries $\Lambda$. |
| $\Delta_\Gamma$ | Diagonal matrix with diagonal entries $\Gamma$. |
| $\Delta_\Phi$ | Diagonal matrix with diagonal entries $\Phi$. |
| $\Delta_\Psi$ | Diagonal matrix with diagonal entries $\Psi$. |
| $\Delta_{\Phi_j}$ | Diagonal matrix with diagonal entries $\Phi_j$ of subject $j$. |
| $\Delta_{\Psi_j}$ | Diagonal matrix with diagonal entries $\Psi_j$ of subject $j$. |

The above joint diagonalization gives us the pair of diagonal matrices Δ_Φ_ and Δ_Ψ_, where the diagonals Φ = {*ϕ*_1_, *ϕ*_2_, …, *ϕ*_*n*_ } and Ψ = {*ψ*_1_, *ψ*_2_, …, *ψ*_*n*_ } constitute the joint eigen spectra of ***L*** and ***F*** respectively. We additionally assume 𝒜 is orthogonal matrix such that 𝒜^*T*^ *A* = ***I***. From the perspective of projection theory, the eigen spectra Φ and Ψ could be viewed as the projections of structural Laplacian and functional connectome on the space spanned by {***a***_1_, ***a***_2_, …, ***a***_*n*_}. For example, *ϕ*_1_ and *ψ*_1_ are the projections of ***L*** and ***F*** respectively on ***a***_1_. Thus, the problem in (1) could be equivalently formulated as a minimization as follows:

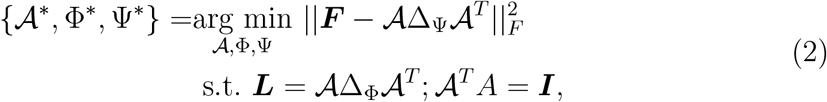

where the subscript *F* stands for Frobenius norm of a matrix.

The mathematical problem in (2) is a particular instance of 𝒜 simultaneous diagonalization. There exist several algorithms to solve (2); here we choose the method by [40] due to its computational simplicity. We refer the columns of 𝒜 as joint eigenmodes of the pair (***L, F***). One attractive aspect of joint eigenmodes is that we can diagonalize both structural (Laplacian) and functional connectome using the same set of these eigenmodes. We note that there could exist an interesting relationship between these joint eigenmodes and individual eigenmodes of structural Laplacian or functional connectome. We discuss this in details in Section 2.2. Suppose (***U***, Δ_Λ_) are the eigenmode and eigen spectrum respectively of structural Laplacian ***L*** and (***V***, Δ_Γ_) are the eigenmode and eigen spectrum respectively of functional connectome. Then,

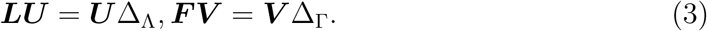

We note that the dominant eigenmodes of structural Laplacian ***L*** are those eigenmodes with least eigen spectrum, whereas dominant eigenmodes of functional connectome ***F*** are those eigenmodes with highest eigen spectrum.

Suppose the eigenvalues of ***L*** are arranged as follows: *λ*_1_ ≤ *λ*_2_ ≤ … ≤ *λ*_*n*_. Then one can show that 0 ≤*λ*_*i*_≤ 2. A detailed proof is quite straightforward by using two facts: (i) symmetric nature of structural connnectome *S* and (ii) non-negativeness of each entry of *S*. We also note that functional connectomes are positive semi-definite matrices because they arise from covariance matrices. Therefore, the eigenvalues in (3) of each functional connectome follows: *γ*_*i*_ ≥ 0, ∀*i*, where *γ*_*i*_ are the diagonal entries of Δ_Γ_.

It remains an open question to explore the connections between the joint eigen spectra and individual eigen spectra. We state two interesting theorems on the bounding properties of joint eigen spectra Φ and Ψ as follows:

### Theorem 1

The joint eigenvalues of structural Laplacian {*ϕ*_1_, *ϕ*_2_, …, *ϕ*_*n*_} are bounded:

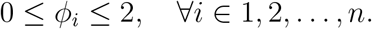

**Proof** : See Appendix 6.1.

### Theorem 2

The joint eigenvalues of functional connectome {*ψ*_1_, *ψ*_2_, …, *ψ*_*n*_} are non-negative.

**Proof** : See Appendix 6.2.

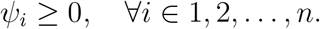

In Figure 1, we display a pair of structural and functional connectomes and their respective eigen spectra (ordered) of a representative subject. It can be seen that the eigenvalues of ***L*** lie in the range [0, 2] and the eigenvalues of the functional connectome are non-negative. The structural and functional eigenmodes corresponding to the dominant/top four eigenvalues ***L*** and ***F*** respectively are shown in second and third rows of Figure 1. The bottom two rows show the joint eigen spectra and top four joint eigenmodes as a result of joint optimization of the ***L*** and ***F*** pair. Notice some visual similarities between the joint eigenmodes with the respective dominant (individual) eigenmodes of structural and functional connectomes. We investigate this property in detail in Section 2.2.

**Figure 1:**
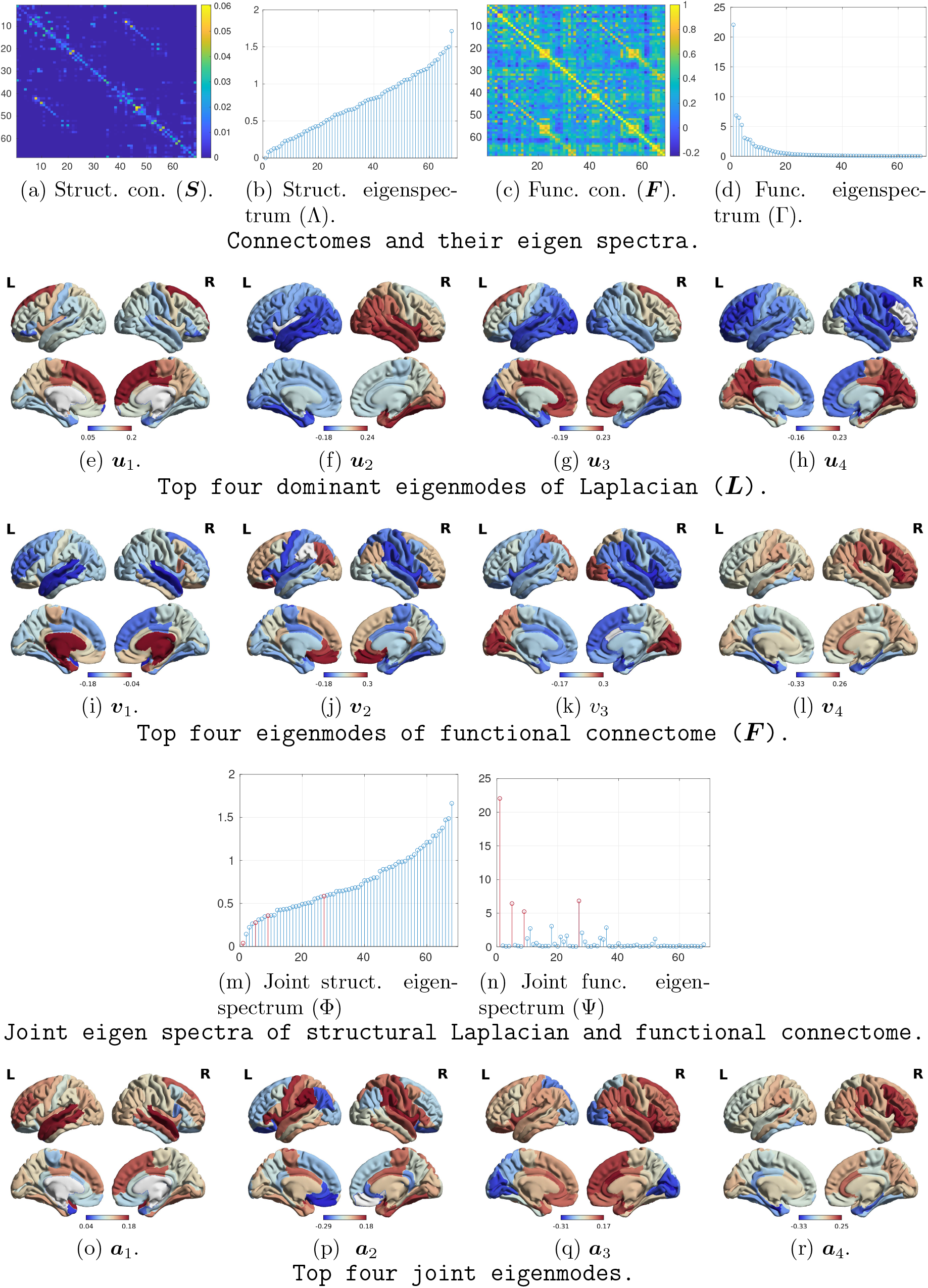
A pair of structural and functional connectomes (from public dataset [39] subject # 1) and their respective eigen spectra. The line plot in (b) shows eigen spectra of Laplacian of the structural connectome shown in (a). Similarly, the line plot in (d) displays eigen spectra of the functional connectome in (c). In second row, we show the top four eigenmodes (eigenvectors corresponding to least eigenvalues) of the structural Laplacian. Similarly, in the third row we show the top eigenmodes of functional connectome. The fourth row show the joint eigenspectra for the structural Laplacian and functional connectome. The bottom row shows the top four dominant (with respect to Ψ) joint eigenmodes.

Figure 2 shows (Φ, Ψ) eigen spectra for eight other representative subjects from [39]. The joint diagonalization is performed independently on (***L, F***) pair for each subject. Visual results from 8 subjects are shown in figure 2. Across all subjects, the first four joint modes lie in the lower-half of both the structural and functional eigenspectra. Also eigenvalues in Ψ for each subject *sparse* in nature, most of the eigenvalues are close to zero except a few. This unique nature of the Ψ motivates us to establish a subspace relationship between vector space of structural Laplacian connectome and vector space of functional connectome of the same subject. We also note that the most dominant eigenmode is often the one corresponds to the least value of Φ. However, subsequent (after the top one) dominant joint eigenmodes do not follow any consistent sequence across subjects.

**Figure 2:**
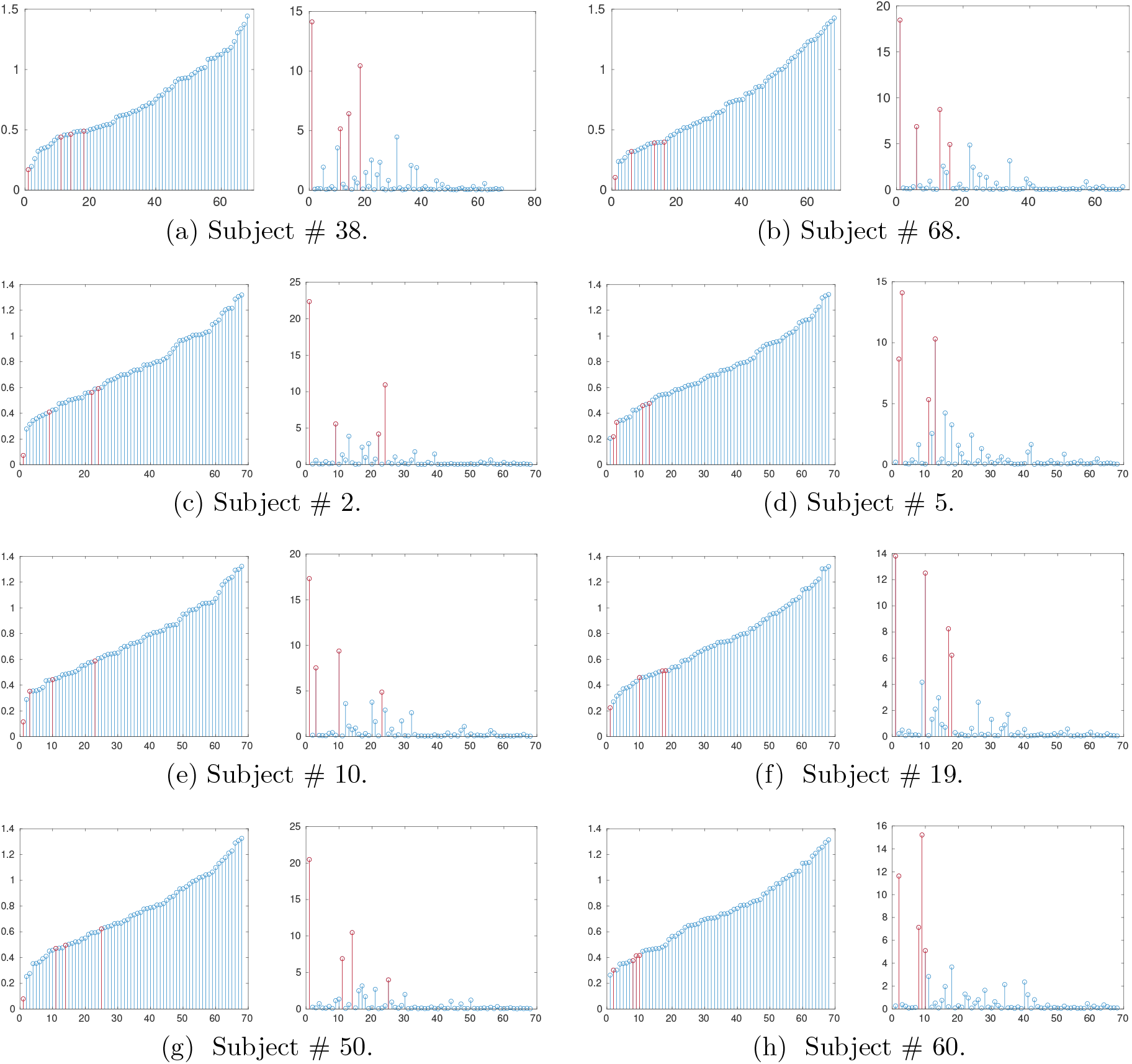
Joint eigen spectra for different representative subjects from dataset [39]. For each subject, left plot is joint eigenspectrum (Φ) of structural Laplacian and right plot is joint eigenspectrum (Ψ) of functional connectome. Across all subjects, it can be noted that the first four joint modes lie in the lower-half of both the structural and functional eigenspectra. However, there is no consistent ordering in the dominant eigenmodes with reference to ascending ordering of joint eigen spectra (Φ) of structural Laplacian.

### 2.1 Subspace Relationship

In this section, we establish the fact that the functional connectome can be reconstructed using only a small number of the joint eigenmodes - i.e. a low-rank subspace. As an example, we re-arrange the joint eigenmodes {***a***_*i*_} ∀*i* ∈ 1, 2, …, *n* and Ψ of the subject from Figure 1(n) in ascending order of the latter and shown in Figure 3(a). We plot Pearson R metric [19] correlation between ground-truth functional connectome and the estimated one from the reduced eigenmodes as a function of number of joint modes (*k*). Here *k* = 1 implies the estimated functional connectome from only one joint mode and *k* = *n* is when all modes are used in the estimation which results in perfect recovery.

**Figure 3:**
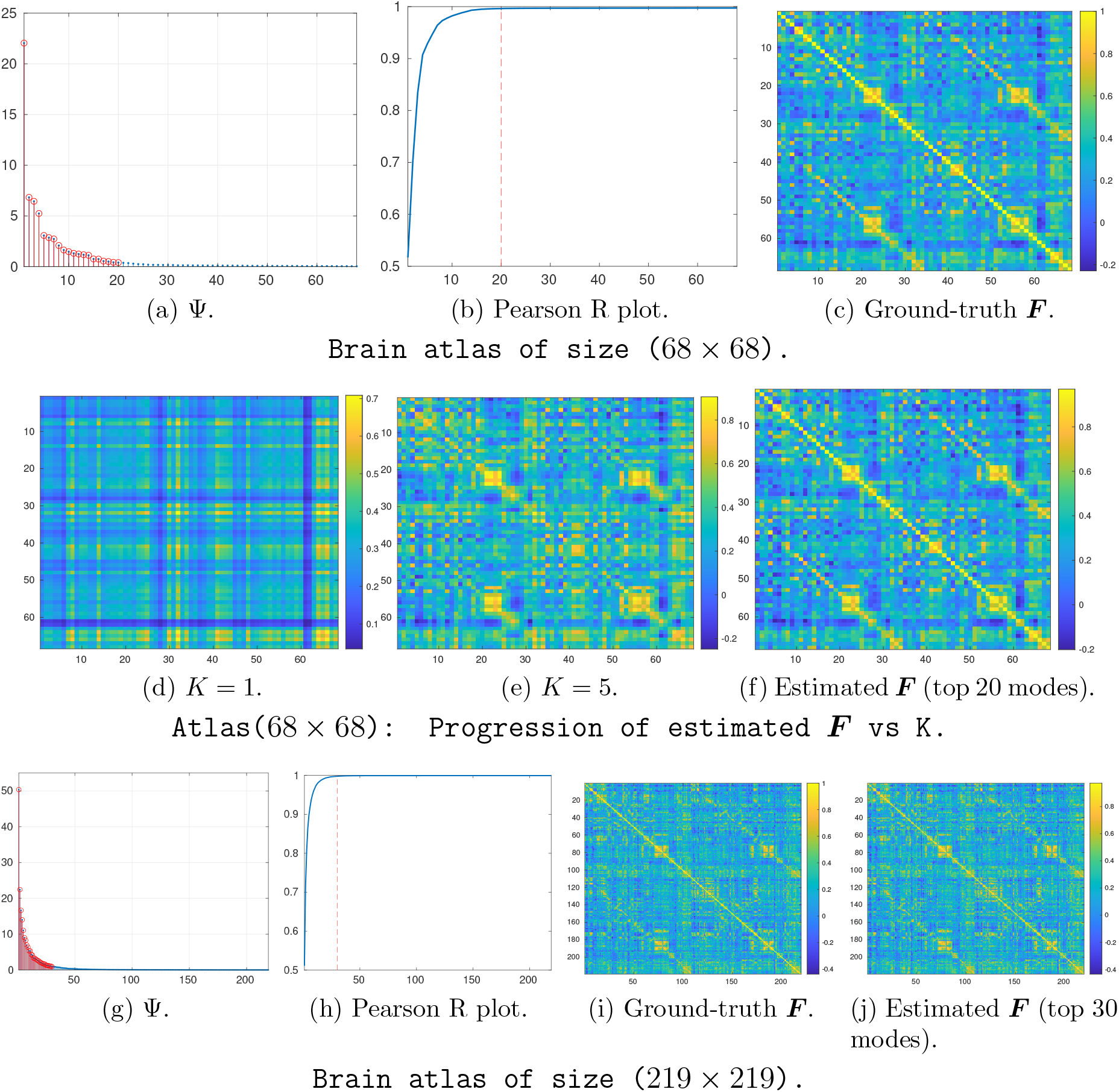
Example of subspace relationship in structure-function joint eigen-spectrum. (a): Ψ obtained via joint diagonalization. Top K = 20 values are marked in red. (b): Plot of Pearson R value as a function of *K*. For each *K*, we estimate functional connectome as in (2.1). The red dotted x-line indicates the instance when estimated functional connectome almost matches with the ground-truth one. In the top example, it is *K* = 20 where the Pearson R is 0.996. Similarly, for *K* = 30, the Pearson R is 0.998 in case of (219 *×*219) atlas in the bottom example. The respective estimated functional connectomes are displayed in the last column. It is evident that both the estimated connectomes are visually indistinguishable to the true functional connectomes shown in third column. These examples demonstrate that functional connectome is contained within structural connectome.

Using the properties of singular value decomposition (SVD), the structural Laplacian can be expressed as outer-products of the joint eigenmodes:

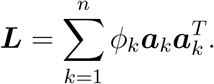

Similarly, the functional connectome can be expressed as the outer-products of the joint eigenmodes. The Pearson R plot in Figure 3(b) suggests that *K* = 20 is enough to approximate ***F***. In particular:

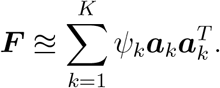

Only a few number (*K*≪ *n*) of joint eigenmodes sufficiently span the functional connectome. Another example is shown the last row in Figure 3. Here only *K* = 30 dominant modes span the functional connectome (*n* = 219). In summary, the existence of joint eigenmodes and the *sparse* nature of Ψ lead to this subspace relationship. Note that the low-rank nature of functional connectome is also preserved in form of joint subspace [17, 19].

### 2.2. Relationship between Joint Eigenmodes and Individual Eigenmodes

Here, we study the relationship between joint eigenmodes and individual eigenmodes of structural Laplacian and the functional connectivity matrix. In particular, we perform quantitative analysis in terms of Pearson R metric [19]. We note that a high value of Pearson R implies high correlation. In Figure 4, we show the results for subject #1 from dataset [39]. For completeness, we also show the correlation with group-level joint eigenmodes of all 68 subjects in [39]. We found a high correlation between the pair (***a***_1_, ***v***_1_) when we compare with individual joint eigenmodes. On the other hand, the group level mode ***a***_2_ is found to have high correction with the functional mode ***v***_2_.

**Figure 4:**
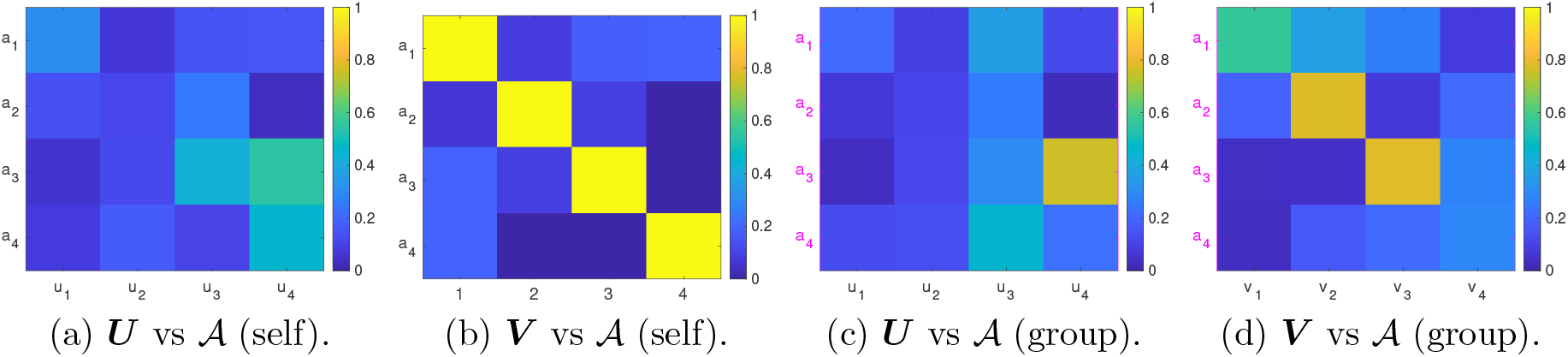
Similarity of joint eigenmodes with structural and functional eigenmodes. Pearson R for different combination of (first four) eigenmodes for subject #1 in dataset [39]: (a) eigenmodes of structural Laplacian of subject #1 vs its joint eigenmodes, (b) eigenmodes of functional connectome of subject #1 vs its joint eigenmodes, (c) eigenmodes of structural Laplacian of subject #1 vs the group-level joint eigenmodes, and (d) eigenmodes of functional connectome of subject #1 vs the group-level joint eigenmodes. A higher value indicates high correlation. With reference to *self* joint eigenmode of this subject, we observe highest correlation between (***v***_4_, ***a***_4_) pair in (b). For the group-level joint eigenmodes, the highest correlation is found between (***v***_2_, ***a***_2_). In general however, the joint eigenmodes are more similar to the functional eigenmodes than the structural eigenmodes at both the individual and the group level.

We recall,

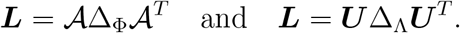

We note that eigenmodes of a real symmetric matrix are orthogonal [43] and span the entire space ℝ^*n*^. In other words, the vector space spanned by the columns of 𝒜, referred as ℂ_𝒜_ has dimensionality *n*. Similarly, the vector space spanned by the column of ***U***, referred as ℂ_***U***_ has dimensionality *n*. Therefore, there exists an invertable mapping **R**_***L***_ : ℂ_***U***_ → *ℂ*_𝒜_, such that:

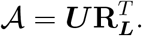

Suppose the vector space spanned by the column of ***V*** is given by ℂ_***V***_, which has dimensionality *n*. Similar to the case of structural Laplacian, there exists a mapping **R**_***F***_ : ℂ_***V***_ → *ℂ*_𝒜_, such that:

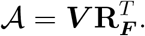

Therefore, the joint eigenmodes and the eigenmodes of the structural Laplacian and functional connectome are all related through rotational or other invertible similarity transformations.

## 3. Structure-Function Mapping

In this section, we extend the concept of joint eigenspace to estimate functional connectivity of a subject from its structural (via Laplacian) counterpart. Given the joint eigenmodes between the structural Laplacian and functional connectome, the task boils down to estimating Ψ from Φ at an individual subject level. It was reported in [19] that there could be an inverse relationship between log(Ψ) and Φ based on a linear graph model predicting a subject’s functional connectivity matrix from their structural connectivity matrix via graph diffusion [17]. Motivated by the findings in [19], here we aim to estimate Ψ from Φ.

### 3.1. Joint Eigen Spectrum Mapping

We propose a data-driven approach to learn a group level mapping between Φ and Ψ. The idea is to consider the diagonals of both Δ_Φ_ and Δ_Ψ_ as two points in a vector space of size ℝ^*n×*1^. Then we pose the mapping between Φ and Ψ as a leastsquares problem [44]. Suppose there are *M* subjects in the group from which learn the least-squares mappings. Therefore, for each subject *j* ∈ {1, 2, …, *m*} there is a pair of joint eigenvalues (Φ_*j*_, Ψ_*j*_) obtained by via performing joint diagonalization of the pair (***L***_*j*_, ***F***_*j*_). First, we embed joint eigenvalues of each subject in two matrices *X* ∈ *ℝ*^*n×m*^ and *Y* ∈ *ℝ*^*n×m*^ as follows:

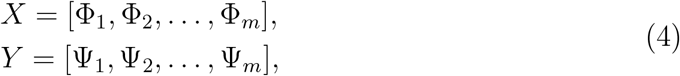

where Φ_*j*_, Ψ_*j*_ ∈ *ℝ*^*n×*1^, ∀*j* ∈ {1, 2, …, *m*}. We consider the following linear (/directional) transformation *W* ∈ *ℝ*^*n×n*^:

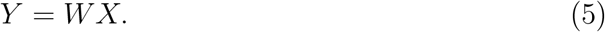

To circumvent the limitation of over-fitting incurred by simple least-squares solution, we seek to impose rank constraint [45] on *W*. Note that this problem is a NP-hard. Various approaches have been proposed in the literature to solve this problem approximately [46]. One efficient approach is to regularize the objective by trace norm (sum of singular values), which is popularly called as nuclear norm. In particular, we consider the following minimization:

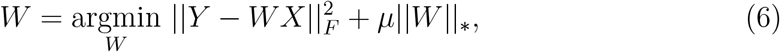

where *μ >* 0 and ||*W* ||_*_ is the nuclear norm of *W*. We solve the minimization using a primal gradient method [45].

The reason we used structural Laplacian instead of structural connectome is as follows. The eigen spectrum of structural Laplacian is bounded in [0, 2]; whereas no such bound exists for the eigen spectrum of structural connectome. We also note it can be shown theoretically that the eigen spectrum of functional connectome is non-negative. Therefore, while working with structural Laplacian, the spectrum mapping W does not exhibit polarity shift. However, in theory, one could simply work directly with structural connectome. In that case, the respective joint eigenmodes would be between structural and functional connectomes.

### 3.2. Predicting Functional Connectome with Joint Mapping

Here we aim to estimate the functional connectome 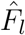 for an individual subject *l*, given their structural Laplacian *L*_*l*_, group level joint eigenmodes and individual joint eigen spectra. An optimal spectrum mapping in (6) would work best under the assumption that joint eigenmodes for each subject is known to us. However, this is not really a realistic assumption from the perspective of a predictive model. We empirically notice that each subject has different set of joint eigenmodes. This is what makes the above formulation as bottleneck to being an efficient predictive model from structural to functional connectome. Instead, our aim is to find a set of group-level joint eigenmodes 𝒜 ∈ ℝ^*n×n*^, where each column represents group-level joint eigenmode. In particular, we consider the following optimization problem:

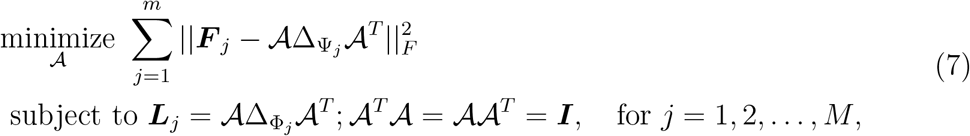

where ***F***_*j*_ and ***L***_*j*_ stand for functional connectome and structural Laplacian of subject *j* respectively. Using eigen spectrum mapping from (5), we get Ψ_*j*_ = *W* Φ_*j*_ which implicitly takes into account the fact that 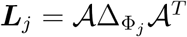. Further, by using standard trace equality and the connection between Frobenius norm and trace of a matrix, we further simplify the objective function above as follows:

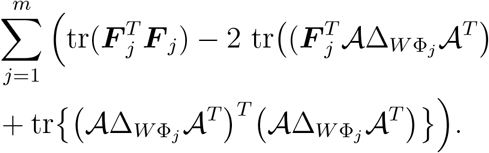

By using *𝒜*^*T*^ *𝒜* = ***I*** and cyclic property of trace of matrix, the third term above is simplified to 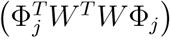. In fact, it is clear that both first and third terms above do not depend on *A*. Thus, the reduced optimization is:

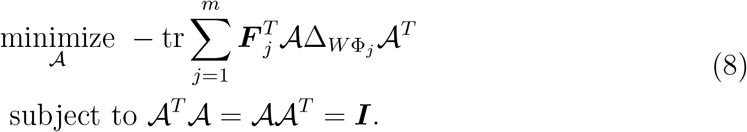

It is worth noting that there is no known closed-form solution to (8). We adopt the iterative algorithm described in [47] to find an approximate solution.

In summary, we propose a two-step strategy for the structure function mapping. First, we construct matrices *X* and *Y* using both structure and function connectomes of the subjects; then we learn the spectrum mapping *W* in (6). Second, we solve (8) to obtain group level joint eigenmodes *A*. In the next iteration, we re-train the mapping *W* in (5) using these joint eigenmodes followed by estimating a refined group joint eigenmodes using the new mapping. Upon convergence, we obtain a final set of joint eigenmodes 𝒜 and spectrum mapping *W*.

Detailed steps of the proposed JESM Algorithm are described in 1. We finally obtain the group level joint eigenmodes 𝒜 and mapping *W*, and estimate functional connectome for an arbitrary subject *l* as follows:

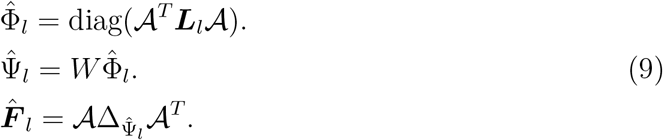

### 3.3. Dataset

To validate the proposed brain connectivity analysis framework, we experimented on data from 68 healthy subjects [39]. The brain data acquisition comprised of (i) a magnetization-prepared rapid acquisition gradient echo (MPRAGE) sequence sensitive to white/gray matter contrast, (ii) a DSI sequence (128 diffusion-weighted volumes), and (iii) a gradient echo EPI sequence sensitive to BOLD contrast. These data was pre-processed using the Connectome Mapper pipeline [48]. Gray and white matter were segmented using Freesurfer and parcelled into total of 68 cortical regions. Structural connectivity matrices are estimated for each subject using deterministic streamline tractography on reconstructed DSI data [49]. Functional data were estimated using the protocol in [50]. This includes regression of white matter, cerebrospinal fluid, motion deblurring and lowpass filtering of BOLD signal.

#### Algorithm 1

Joint Eigen Spectrum Mapping (JESM)

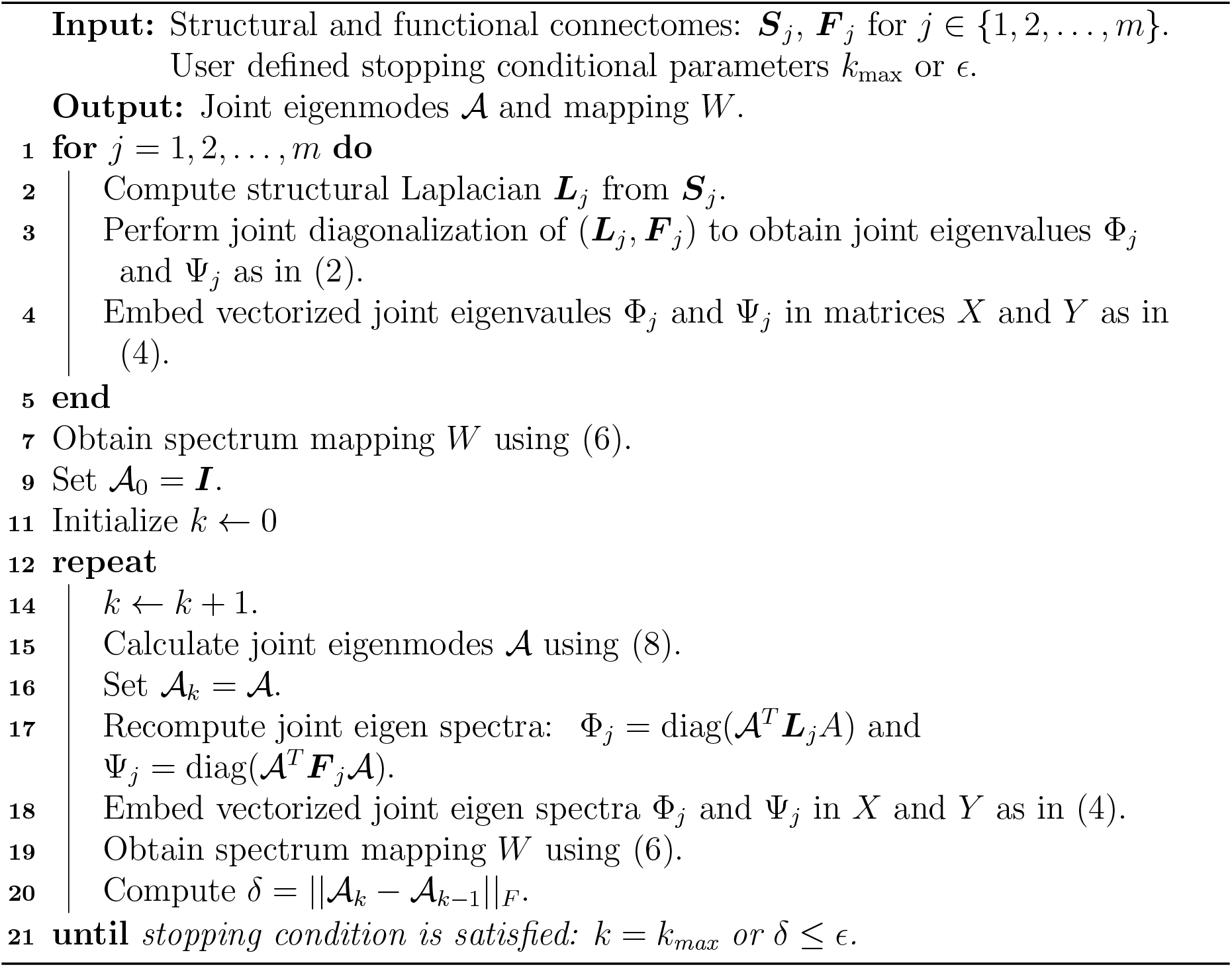

In addition, we have used a schizophrenia dataset [51] consisting 27 control subjects and 27 schizophrenia subjects. All connectomes in this schizophrenia dataset have 83 ROIs; of which 68 cortical and 15 sub-cortical regions.

### 3.4. Benchmark Comparisons

We compare the performance of JESM with the following benchmark methods in terms of estimating functional connectivity of a subject from its structural counterpart.

1. Abdelnour et al. [19]: The standalone eigenmodes of structural Laplacian Λ were used to estimate the functional connectome of the subject. Following [17], the functional eigen spectra Γ were estimated using an exponential transformation of Λ.
2. Tewarie et al. [33]: We implemented the optimization framework of [33] to estimate the projections of functional connectome on the eigenmodes of structural connectome. Note that this work did not focus on a predictive pipeline. Therefore it did not include any form of spectrum or curve fitting. The core focus was on the theory to approximate functional connectome of a subject by the projection of each eigenmode of structural connectome. It was theoretically shown that given the eigenmodes of the structural connectome, the best of estimation of the functional connectome is eigen spectral composition of the projection of functional connectome on each eigenmode. In our implementation, we display the best predicted functional connectome result which could be obtained when one is somehow provided with the optimized projection values.
3. Becker et al. [30]: We implemented the structure-function mapping method in [30], where given both structural and functional connectomes of a particular subject, a rotation matrix and a polynomial expansion of the respective eigen spectra were used to estimate the functional connectome from structure. The core idea is to map the functional connectome from the structural connectome of a subject via a rotation operation. Authors [30] further shown that such a rotation matrix could be derived only if we have access to stand-alone eigenmodes of both functional and structural connectomes for a particular subject. Therefore, it is not straightforward to directly estimate functional connectome from the structural connectome for a subject. However, it could be possible to construct a representative mapping in terms of rotation matrix for a group of subjects via some form of manifold optimization.

In summary, both Abdelnour et al. [19] and Tewarie et al. [33] rely on the eigenmodes of structural connectome (or Laplacian). On the other hand, Becker et al. [30] makes use of structural eigenmodes in an implicit manner to construct the rotation matrix based mapping.

### 3.5. Performance Evaluation of the Predictive Model

We study the effectiveness of the proposed method in terms of accurately estimating functional connectome (FC) of a subject from its structural connectome (or Laplacian) and compare with state-of-the-art methods in the literature. The prediction performance is quantified in terms of Pearson R statistic between the estimated FC and true FC of a subject.

In our group level prediction pipeline, we learn the mapping between joint eigenvalues of structural Laplacian and eigenvalues of functional connectome. We perform 4-fold cross validation experimental setting to evaluate and compare the predictive methods. First, we use both structural and functional connectomes of the train set of subjects. We perform joint diagonalization and then obtain the initial estimate the linear mapping. Next we learn group level joint eigenmodes by solving an optimization problem using the toolbox in [52]. Both the mapping and joint eigenmodes are further refined through few iterations. Finally, we use the learned mapping and group level joint modes to predict functional connectome from the structural connectivity of each subject from the test set.

## 4. Results

In this section, we first display and the group-level joint eigenmodes of a group of 68 healthy subjects in the dataset [39]. In particular, we run Algorithm 1 on the collection of connectome-pairs to obtain the group-level joint eigenmodes. 𝒜 In Figure 5, we display top four the joint eigenmodes, which are the first four columns of 𝒜. We draw few important observations on the the group-level eigenmodes shown. The first mode in Figure 5(a) exhibits relatively low variation across the ROIs. This is perhaps, represent a low frequency mode of the underlined brain graph obtained from the structural Laplacian [3]. The second eigenmode in 5(b) has an interesting distribution across the ROIs. It is found to have prominent intensity gradient from perceptual and motor regions to default mode network (DMN) [53]. Note that a similar intensity gradient profile is also present in the fourth eigenmode. Also notice that the intensity profile in the third eigenmode exhibits two polarity nature across the frontal and peripheral regions in the brain. In Figure 6, we compare the correlation of these four group-level eigenmodes with 7 canonical networks [54]: default, dorsal-attention, frontoparietal, limbic, somatomotor, ventral-attention, and visual. In terms of Pearson R mertic (deep yellow is high), we see that ***a***_2_ joint mode closely resemble with default detwork; ***a***_3_ has positive alignment with both frontoparietal and ventral-attention networks; and ***a***_4_ has high correlation with both dorsal-attention and frontoparietal networks. With reference to geodesic metric, ***a***_1_ mode exhibits high non-Euclidean proximity with all 7 networks and dorsal-attention also has high similarly to all 4 group-level joint eigenmodes.

**Figure 5:**
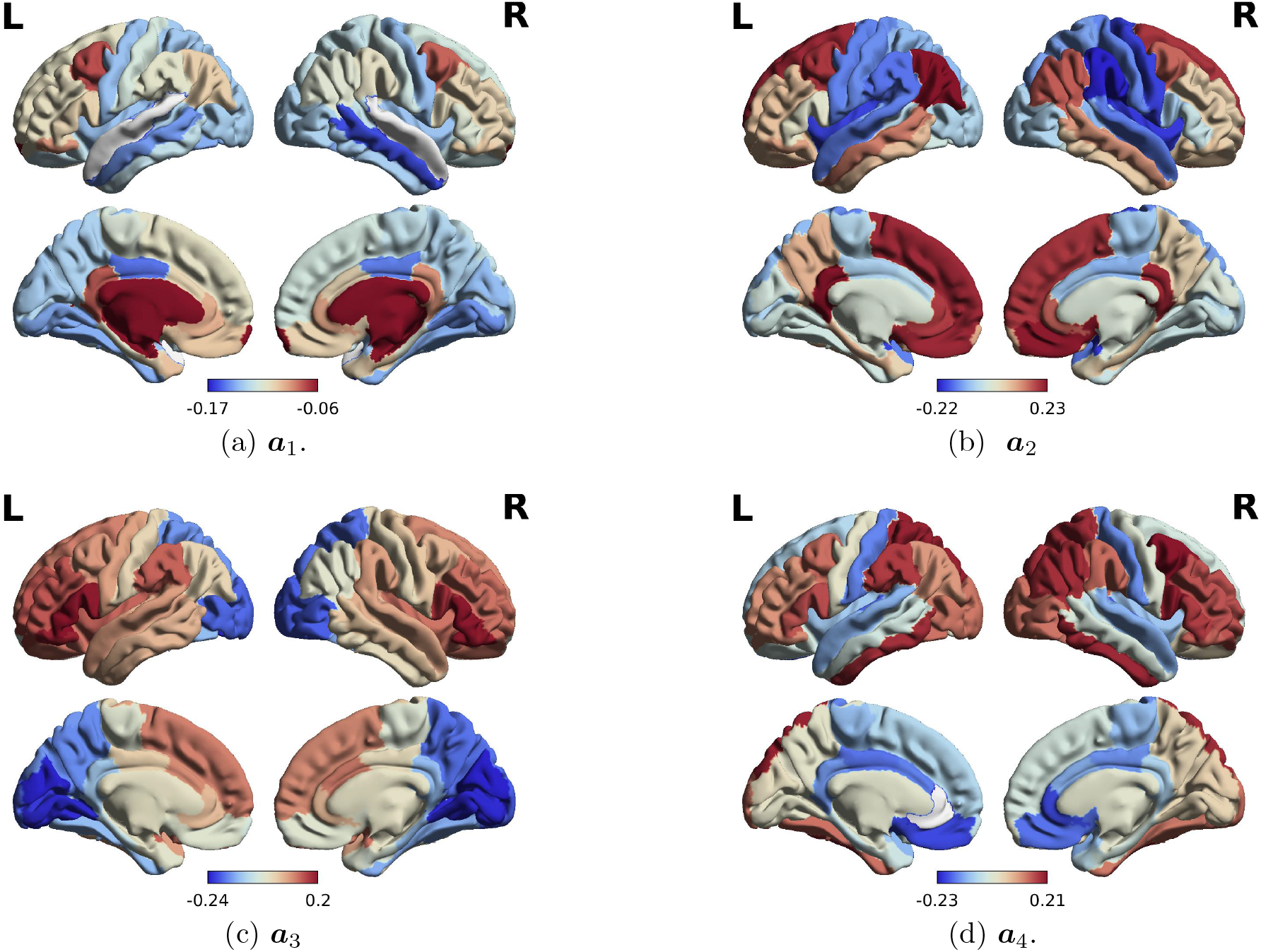
Group-level joint eigenmodes: first four modes. These modes are obtained using Algorithm 1 on all 68 subjects in the public dataset [39].

**Figure 6:**
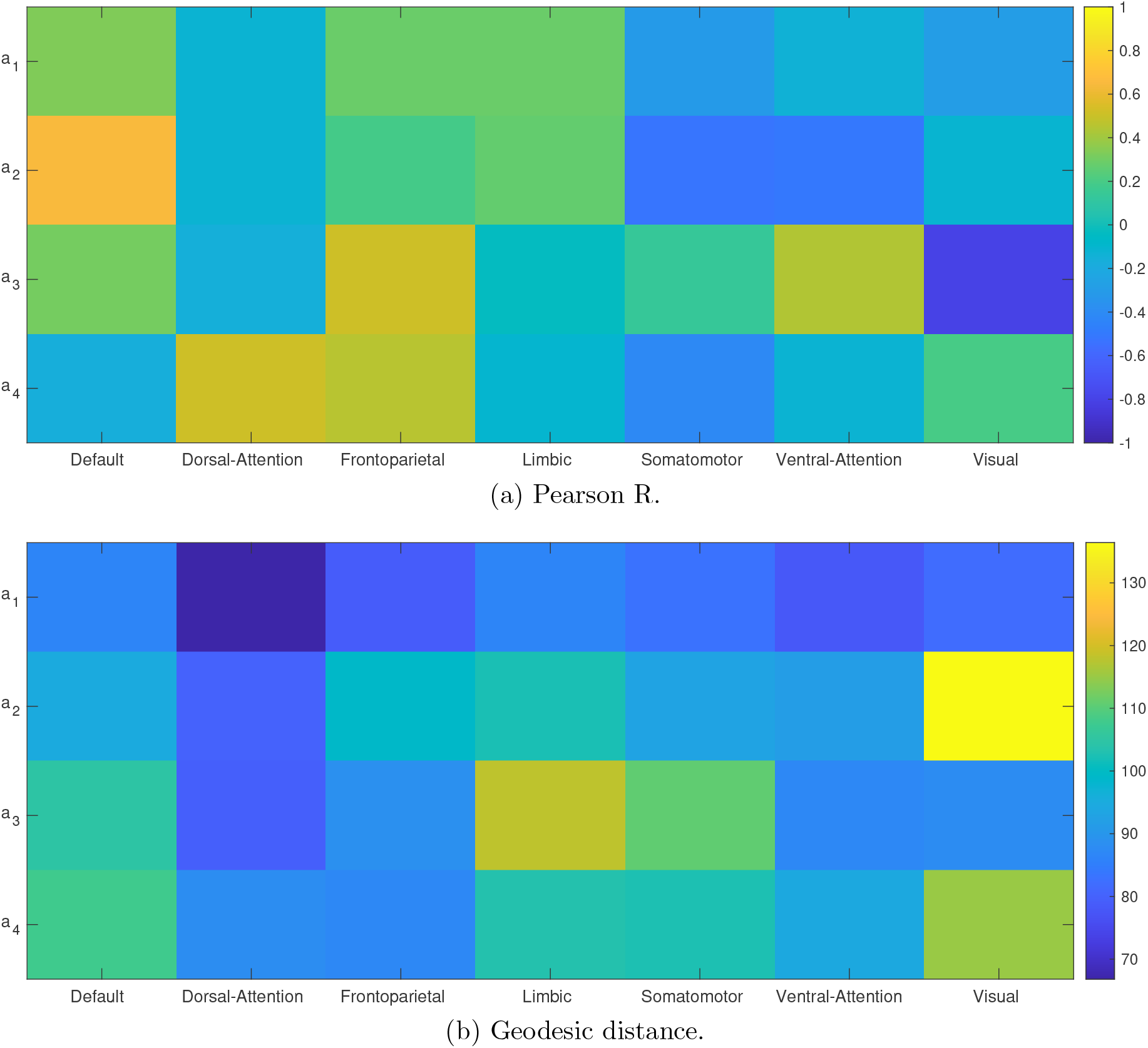
Similarity between 7 canonical networks and group-level (first four) joint eigenmodes of 68 subjects in dataset [39]. The pair (***a***_2_, default) has the highest similarity in terms of Pearson R metric. The pair (***a***_1_, dorsal-attention) has the highest similarity in terms of geodesic metric.

### 4.1. Comparison of Structure-Function Mapping Performance

Next, we perform detailed experiments to examine the effectiveness of the proposed joint eigenmode approach in structure-function mapping. We start with detailed experimentation of our proposed eigen spectrum mapping approach. Finally, we compare with state-of-the-art methods in the literature for structure-function mapping.

In Figure 7 we present results of estimated functional connectomes using our method for two representative subjects [39]. We also display the results for the benchmark methods: Abdelnour et al. [19], Tewarie et al. [33], and Becker et al. [30]. We first group the dataset [39] of 68 healthy subjects into two groups and then perform 4-fold cross validation setting for evaluation of the predictive methods. Note that the connectomes used in this experiment have 68 cortical regions. The respective Pearson R [19] values and geodesic distances [55] are noted below each panel. A higher value of Pearson R and lower value of geodesic distance indicate better prediction. It is visually evident that proposed JESM outperforms both Abdelnour et al. [19] and Tewarie et al. [33]. The estimated functional using JESM more closely resembles the ground-truth functional connectomes respectively. We summarize the comparison of our proposed algorithm and benchmarks for structure function prediction using violin plots for performance metrics across all subjects in Figure 8. For this analysis, we group all 68 subjects into two and perform 4-fold cross-validation on each group containing 34 subjects. Please note that each connectome has 68 ROIs. We used 4 quality metrics - Pearson R, geodesic distance [55], structural similarity index measure (SSIM) [56], and mean squared error (MSE) [44] to quantify the prediction performance. For Pearson R and SSIM, a higher values indicate better prediction. On the other hand, for geodesic distance and MSE, a lower value refers to superiority of the method. We further present a result in Figure 9 on analyzing estimation at (a) intra - vs inter-hemispheric and (b) short vs long connections. In particular, we generate histogram of the residual (upon linear model fitting) between the predicted and ground-truth functional connectomes. Notice that both in cases the residual histogram is zero-centered. This result suggests that functional connections are neither under-nor over-estimated. In other words, the functional connectivity values are well-estimated across ROIs.

**Figure 7:**
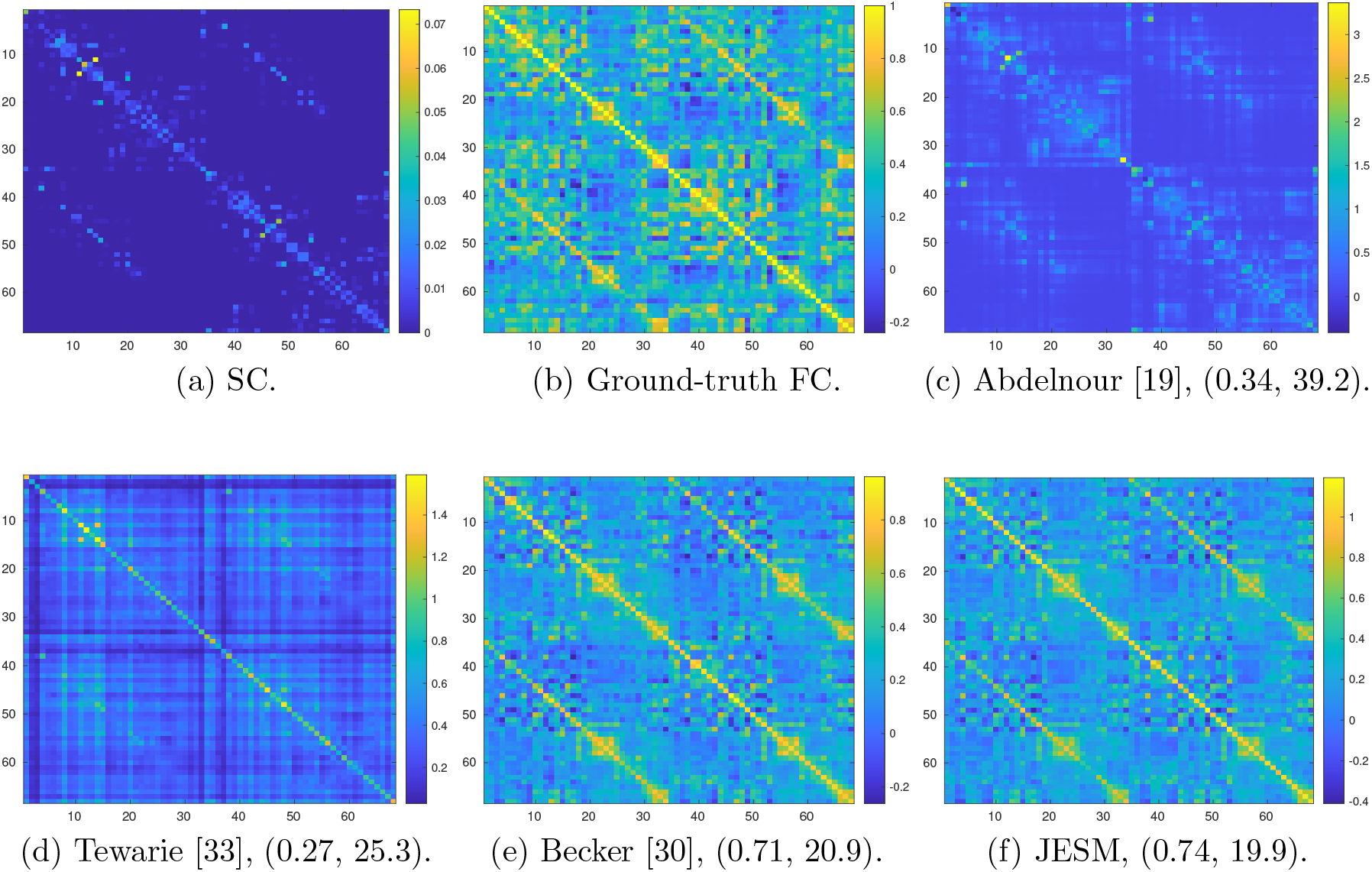
Comparison of structure function mapping for a representative subject (#7). The Pearson R and geodesic distance values for the estimated FC (with reference to the ground-truth one) are reported in the sub-captions. A higher R value indicates superior prediction. On the other hand, a lower value of geodesic distance implies better quality. Among all the methods, our proposed method JESM achieves best results in terms of both visual quality and numerical metrics.

**Figure 8:**
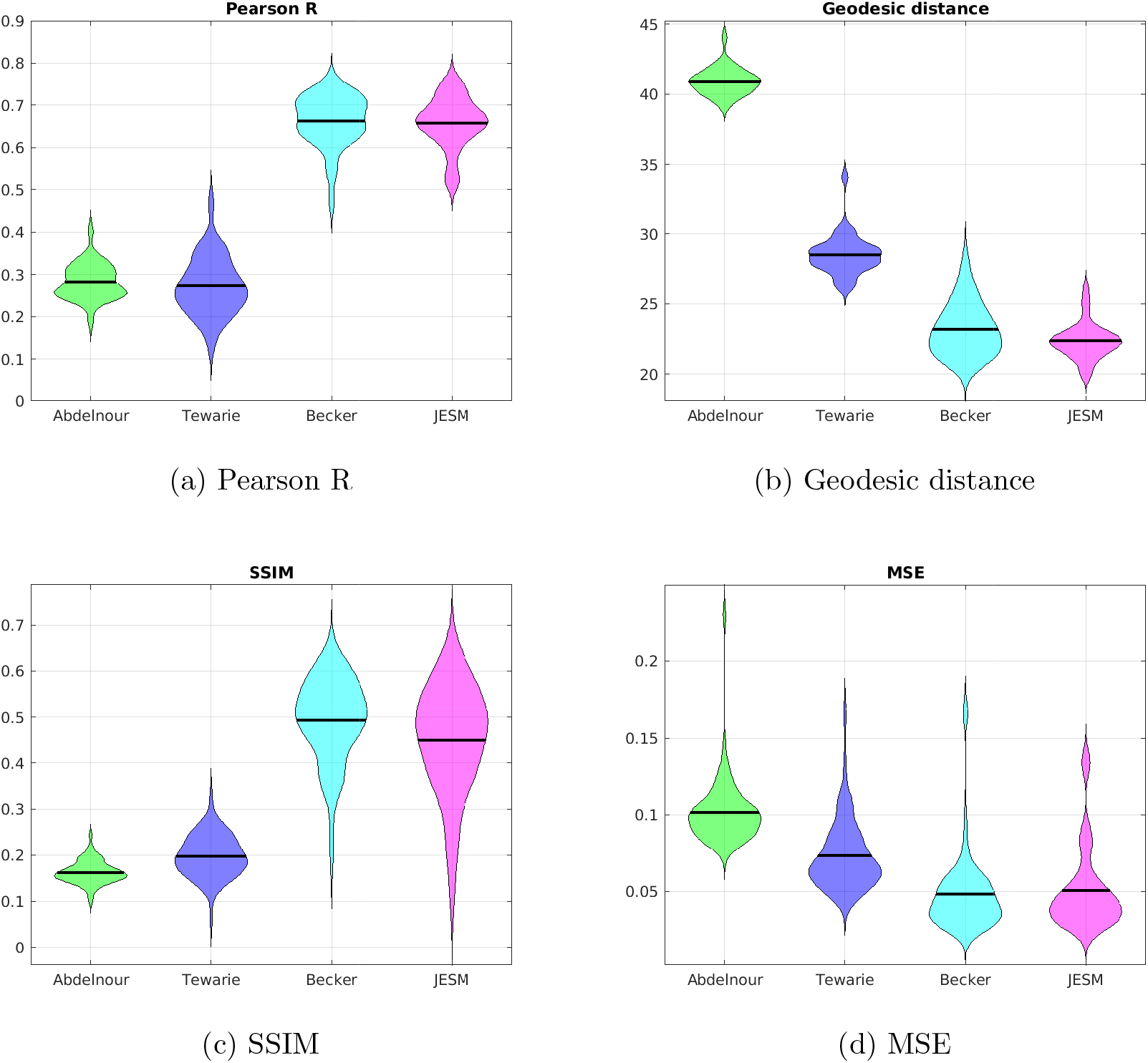
Performance comparisons of structure-function mapping on 68 healthy subjects from [39]. Metrics of performance are: (a) Pearson R, (b) Geodesic distance, (c) Structural similarity index measure (SSIM), and (d) Mean square error (MSE).

**Figure 9:**
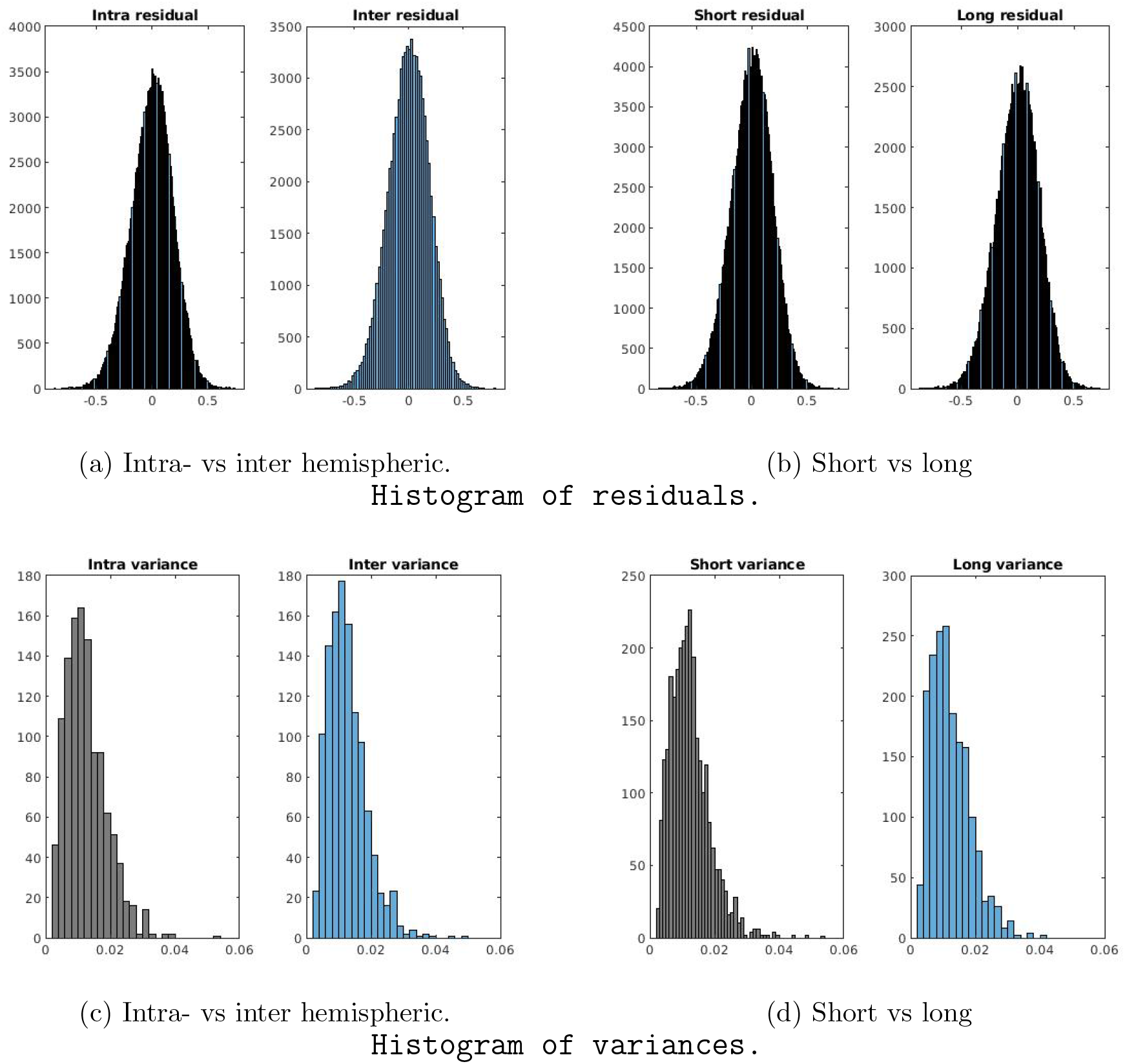
Residual and variance measures on the predicted functional connectivity values for the experimental setting in Figure 8. Notice that the histograms are fairly symmetric distributions with zero mean at (a) intra- vs inter hemispheric and (b) short vs long cases. The mean and skewness of the histograms in (a) are (0, -0.172) and (0,-0.167). The mean and skewness of the histograms of short and long connections in (b) are (0, 0.174) and (0, 0.172). The variance of FC estimates across subjects at intra- and inter-hemispheric regions shown in (c) have the same mean and median values as (0.012, 0.011). Similarly, the variance at both short and long connections shown in (d) have mean and median values (0.013, 0.012). Therefore, JESM preserves the difference across subjects equally well at both intra- vs inter-hemispheric regions as well as short vs long connections in the brain.

We note that our JESM method gives competitive structure-function mapping performance the benchmark methods In fact, we argue that our predictive model could asymptotically match [30] under a suitable rotation of the eigenmodes. However, the important aspect is that our proposed approach has an interesting geometric interpretation. For a given subject, the existence of joint eigenmode could serve as a basis for both structural and functional connectomes. In other words, the joint eigenmode could be used as a mutual representation of the multimodal connectomes. In our current work, we have explored these joint modes to predict functional connectome from structural connectome of a subject. In theory, one could also extend the idea to predict structural connectome from the functional counterpart; which we consider as a future exercise. In a different direction, one could analyze the joint-mode for more than two connectome; for example - fMRI connectome, MEG connectome, and EEG connectome. This could be a new insight to capture the mutual information present in three different functional modalities.

### 4.2. Structure-Function Mapping of Schizophrenia Disorder

In this subsection, we first study the nature of joint eigenmodes of the schizophrenia dataset using our JESM method. In Figure 12, we display first four joint eigenmodes from a set of 27 schizophrenia subjects [51]. Note that the dimension of each connectome in this dataset is 83*×*83; which consists of 68 cortical and 15 sub-cortical ROIs. However, for visualization purposes, we display only the cortical ROIs. Notice the visual difference between joint modes in Figures 5 and 12.

#### 4.2.1. Structure-Function Mapping from Joint modes of Healthy Subjects

To strengthen the analysis of our approach in context group-level joint modes based structure-function prediction, we use the joint modes from an external, completely separate dataset. In this experiment, we train the joint modes and joint eigen spectrum mapping on a set of 27 control subjects from the cohort studied by [51].

Then, we test the prediction (of functional connectome) on Schizophrenia patients who were not present in the training set. Two such visual results are shown in Figure 10. Note that all the connectomes used in this experiment contain 83 ROIs. The predicted functional connectome in (c, f) quite resemble the ground-truth in (b, e) respectively. In Figure 11, we present the performance statistics of the functional connectivity prediction on all 27 Schizophrenia patients and also compare with the benchmark methods. This experiment confirms that the proposed method JESM generalizes beyond healthy subjects.

**Figure 10:**
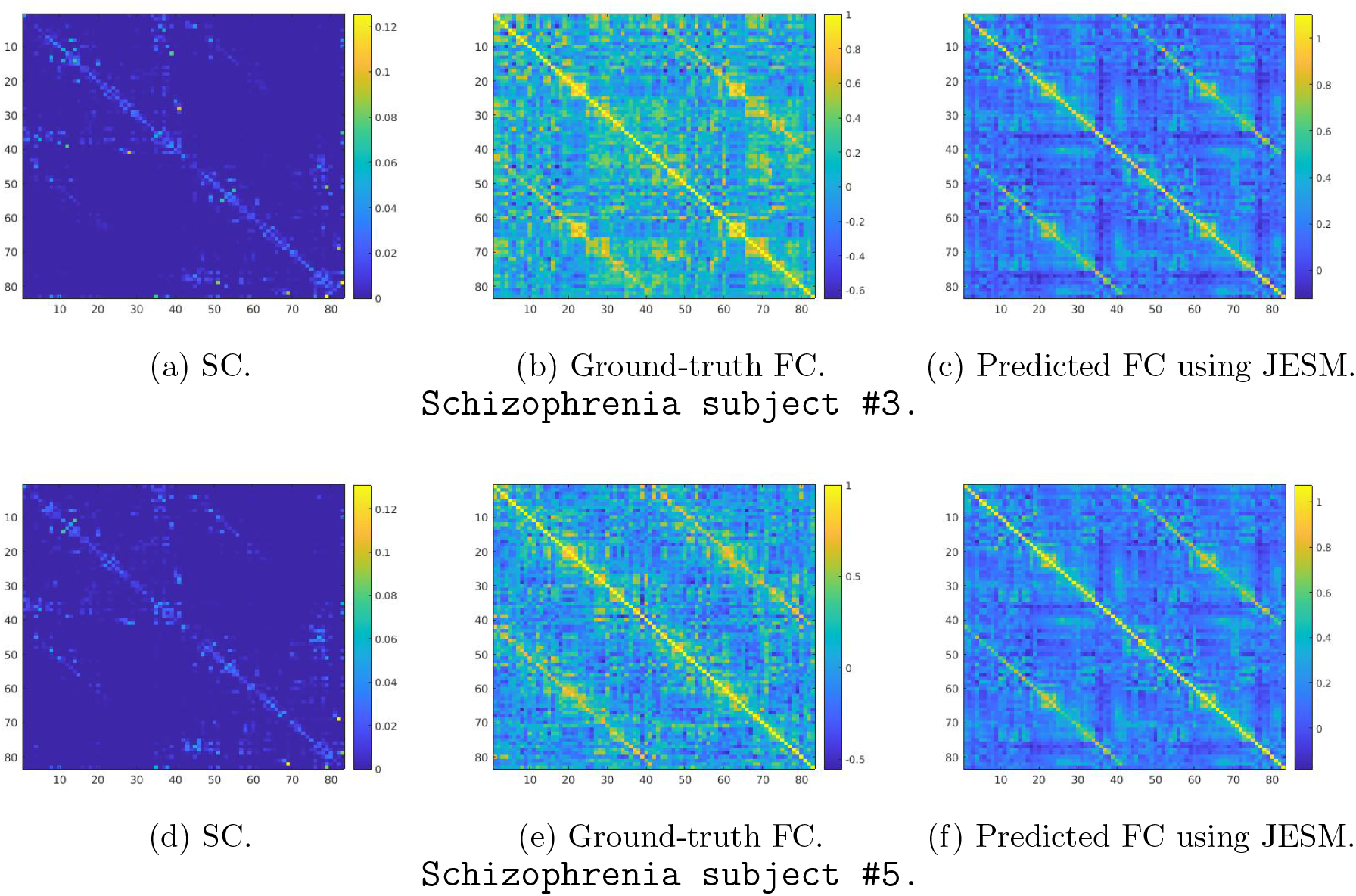
Predicting FC on Schizophrenia dataset [51]: group-level joint modes from healthy subjects. We train the model on connectome pairs of 27 control subjects and then estimate the FC of Schizophrenia patients using our proposed method JESM. The geodesic distance between (b, c) pair is 30.07 and (e, f) pair is 31.67.

**Figure 11:**
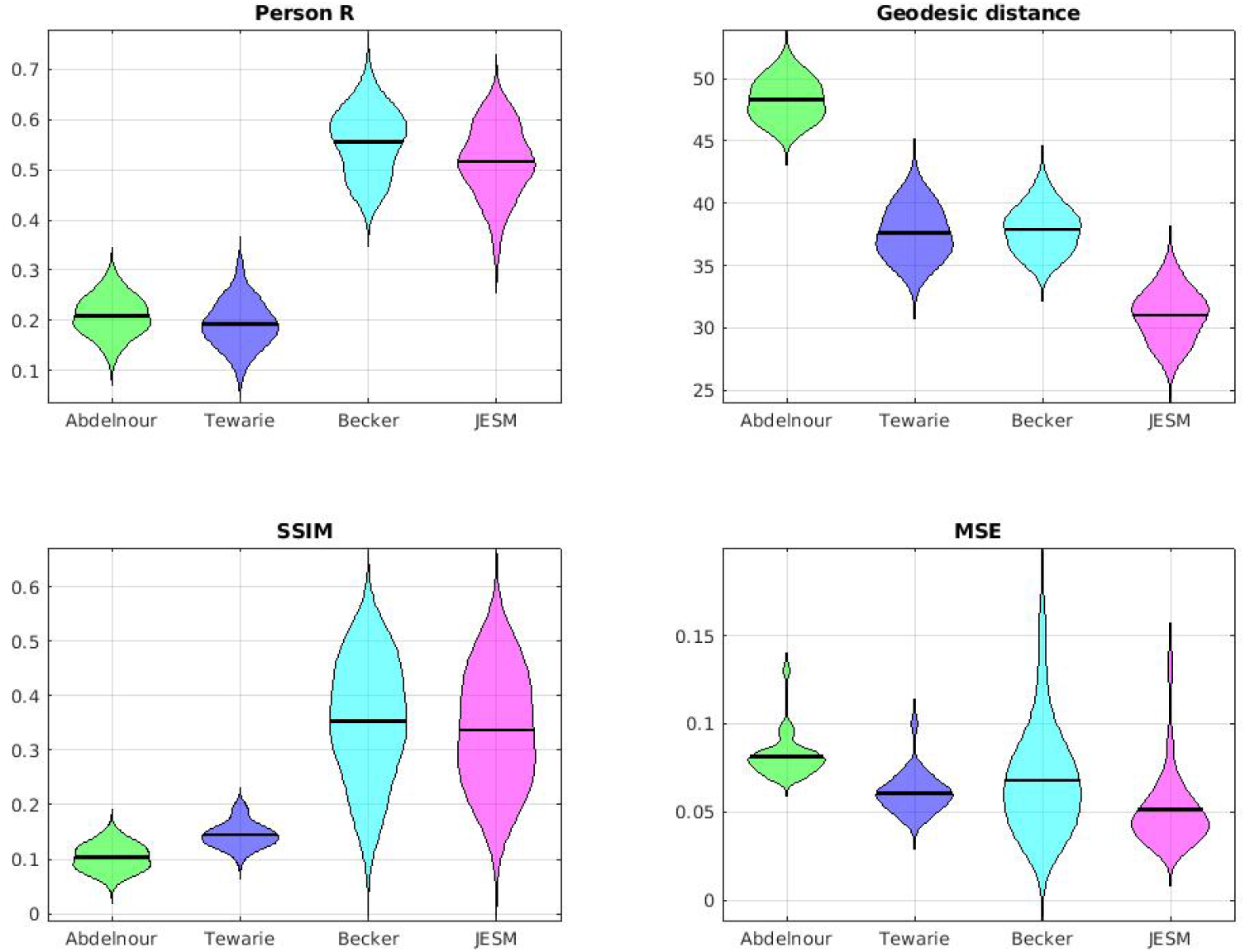
Statistics of the functional connectivity estimation on Schizophrenia patients data [51]: group-level joint modes from healthy subjects.

#### 4.2.2. Structure-Function Mapping from Joint modes of Schizophrenia Subjects

Here study the structure-function mapping performance of JESM using group-level joint modes of Schizophrenia subjects. In particular, we perform 3-fold crossvalidation on 27 schizophrenia subjects. Therefore, in each fold, we learn the joint modes from 18 schizophrenia subjects and use them for the prediction of remaining 9 subjects. In Figure 13, we display a visual result for a subject (from [51] dataset). It is evident that the predicted functional connectome looks very similar to ground-truth. We summarize the comparison of our proposed algorithm and benchmarks for structure-function prediction for schizophrenia data using violin plots in Figure 14. For simplicity, we report only Pearson R and geodesic distance results. The pair of violin plots suggest that our method JESM produces competitive performance in this setting of predicting functional connectome using group-level joint modes of schizophrenia subjects.

**Figure 12:**
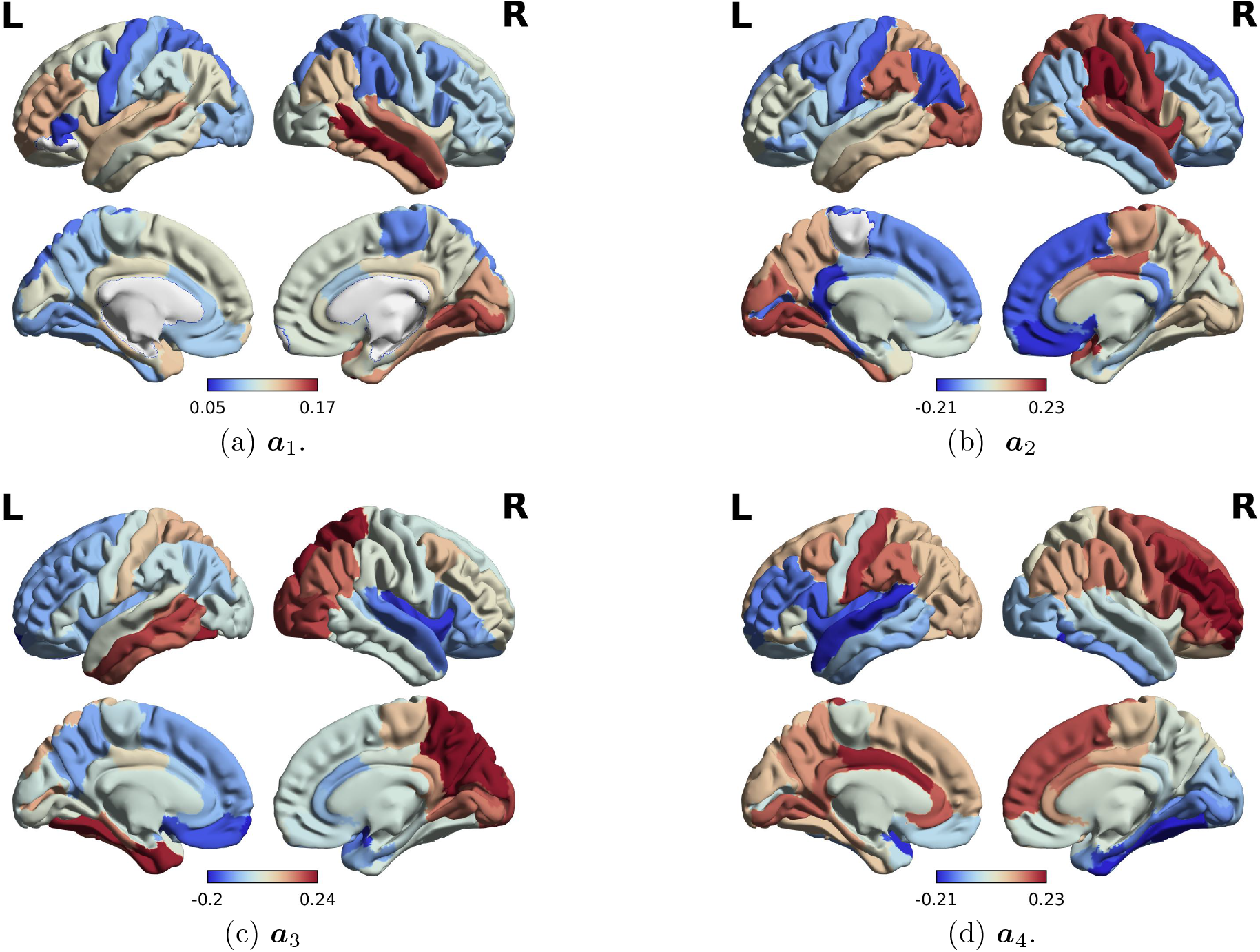
Group-level joint eigenmodes of schizophrenia subjects: first four modes. These modes are obtained using Algorithm 1 on all 27 subjects obtained from the public dataset [51].

**Figure 13:**
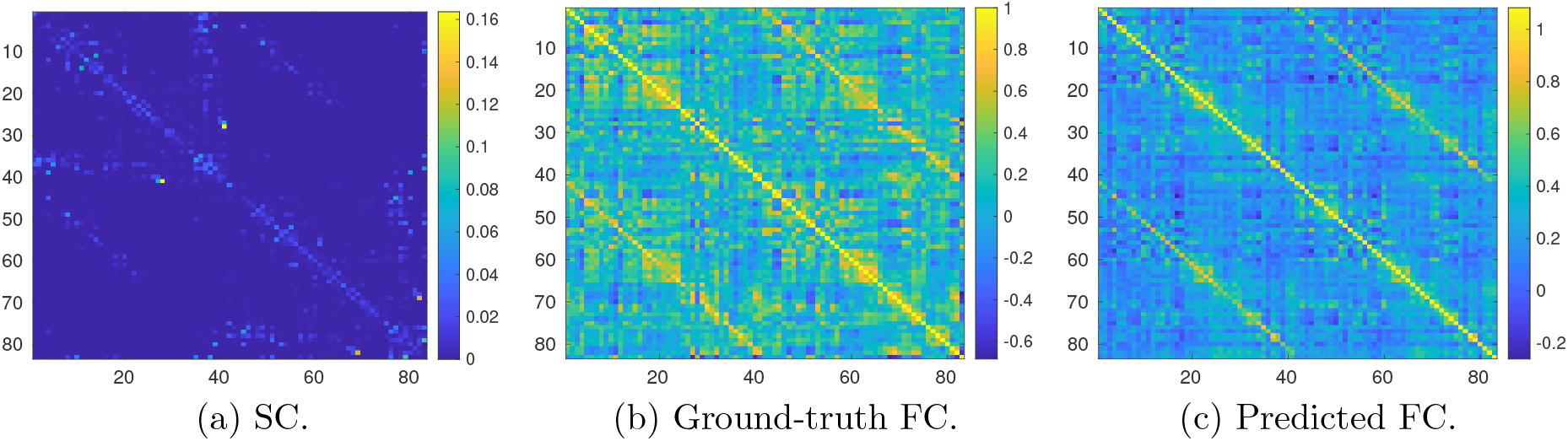
Visual comparison of structure function mapping on Schizophrenia Subject #2. In this experiment, we perform 3-fold cross-validation on 27 schizophrenia subjects. In particular, data from 18 subjects are used to train the group-level joint modes.

**Figure 14:**
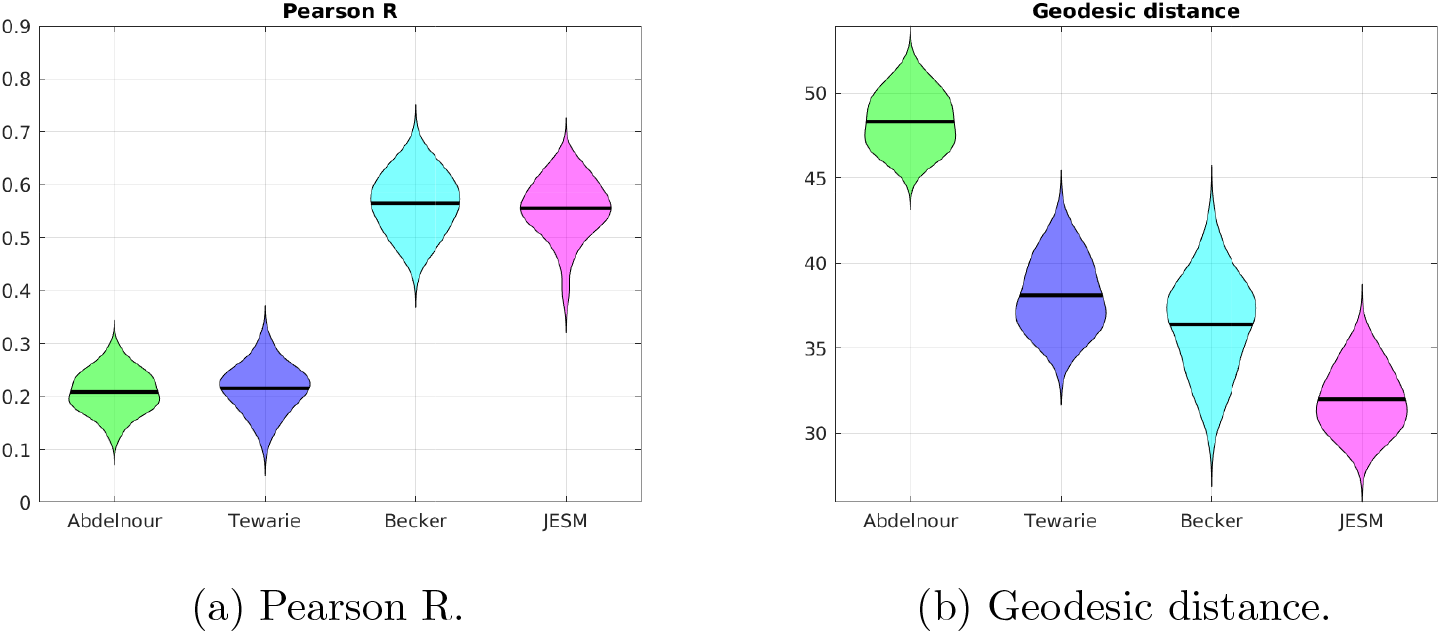
Statistics of the functional connectivity estimation on Schizophrenia dataset [51] with 27 patients. The group-level joint modes are learned from schizophrenia subjects via 3-fold cross validation.

## 5. Discussion

We provided a principled mathematical framework to connect the structural and functional connectivity matrices. By expressing both connectomes using a common eigen space with biophysical interpretability, we simplified subspace relationship between structural and functional connectomes with a joint harmonic mapping between structure and function. We then developed a novel algorithm for predicting functional connectomes based on structural connectomes and this joint mapping, and demonstrate the superiority of this prediction algorithm compared to existing benchmarks. Our theoretical framework generalizes benchmark approaches that predict functional connectomes using a rotation matrix applied to structural eigenmodes and polynomial mapping of structural eigenspectra [30, 36].

The group level predictive model is of particular interest to us. For predicting the functional connectome for a particular subject from its structural connectome via joint subspace mapping, we need to know: (i) the mapping between joint eigen spectra of both and (ii) joint eigenmodes which closely diagonalize both connectomes. We address this concern by two-step optimization. First, we learn a mapping from joint eigenvalues of structural Laplacian to joint eigenvalues of functional connectome via spectrum mapping. Second, we use this learned spectrum mapping to estimate joint eigenmodes using manifold optimization. By efficient use of advanced numerical optimization, we obtain a group level eigenmodes that jointly diagonalize individual subject level structural and functional connectomes.

Among the notable existing works, [19] predicts the functional connectome using the eigenmodes of structural Laplacian. Authors in [33] formulated this structure function mapping as a *L*_2_ minimization problem where the feasible eigenmodes were restricted to the individual eigenmodes of structural connectome. Becker et al. [30] proposed an efficient eigenmode mapping, by introducing a rotation matrix along with the structural eigenmodes as a means to capture individual subjects’ FC. We show below that our methods are computational equivalent barring a few differences, with better interpretability offered by our method in contrast to Becker et al. First, for a single subject, we use a linear projection of the structural and functional eigen spectrum, whereas Becker et al. [30] uses a polynomial approximation. Second, the rotation matrix *R* in [30] can be viewed as somewhat equivalent to our joint eigenmode matrix 𝒜 with a key difference though. Note that 𝒜 jointly diagonalizes both ***F*** and ***L***, whereas *R* does not diagonalize either ***F*** or ***L***. Therefore, our proposal is more geometrically comprehensive with stronger mathematical interpretability. Third, for group-level predictive model, the joint approximation in JESM and Becker et al. [30] are equivalent except for the above differences. Therefore, it is not a surprise that our result is similar to Becker et al. [30].

To study the connection between the structural Laplacian eigenmodes, functional connectome eigenmodes, and joint eigenmodes, we also recall the recent works in [57, 58]. The role of structural eigenmodes in the formation and dissolution of temporally evolving functional brain networks was shown in [59]. Authors in [57] investigated the association between spatially extended structural networks and functional networks using a multivariate statistical technique, partial least squares. They studied whether network-level interactions among neural elements may give rise to global functional patterns. With sufficient experimental results it was demonstrated that the network organization of the cerebral cortex supports the emergence of diverse functional network configurations that often diverge from the underlying anatomical substrate. A multimodal connectomics paradigm utilizing graph matching to measure similarity between structural and functional connectomes was presented in [58]. In this paper, we also performed a statistical analysis to find the similarity in terms of Pearson R metric. We found that the joint eigenmodes are more similar to the functional eigenmodes than the structural eigenmodes at both the individual and the group level. To address the neuroscientific significance of the joint eigenmodes, we compared four group-level eigenmodes with the seven canonical cortical networks: default, dorsal-attention, frontoparietal, limbic, somatomotor, ventral-attention, and visual. Authors in [58] reported high matching between structural and functional connectivity was shown at visual and motor networks are higher as compared to other more integrated systems. It presented an insight into the structural underpinnings of functional deactivation patterns between default mode network (DMN) and task positive systems in brain. Authors in [60] presented a study to systematically analyze the consistency of connectomes, that is the similarity between connectomes in terms of individual connections between brain regions and in terms of overall network topology. The comprehensive study of consistency in connectomes for a single subject examined longitudinally and across a large cohort of subjects cross-sectionally, in structure and function separately. In contrast, our joint eigenmodes are somewhat associated with canonical networks depending on the metric of similarity. Specifically, we found that the pair (a2, default) has the highest similarity in terms of Pearson R metric and the pair (a1, dorsal-attention) has the highest similarity in terms of geodesic distance metric.

Our proposed idea of joint eigenmode decomposition looks promising in terms of both multimodal brain connectivity analysis and structure-function mapping. The main limitation of our estimation approach is computational complexity and numerical stability of group estimates, especially for manifold optimization with a large number of subjects and higher spatial resolution connectomes. Also, in this paper, we only examined the joint optimization of the structural Laplacian and fMRI functional connectomes. The sensitivity of the joint eigenmodes to identify both state and trait characteristics remain to be established. Specifically, explorations of whether joint eigenmodes estimate state changes in task-induced functional data, or trait changes in neurological or psychiatric diseases are needed in future studies.

In summary, our contribution is a step forward in understanding multimodal brain connectivity: underlying structure-function networks that support the differences between diseased and healthy populations. The concept of common subspace in our predictive model could open the door to better data-driven tracking of brain diseases progression. Our long-term vision is to obtain potentially sensitive structure-function biomarkers for differential diagnosis or prognosis or therapeutic monitoring in various neurological disorders.

## 6. Appendix

### Lemma 1

For any ***x*** ∈ ℝ^*n×*1^ spanned by joint eigenvectors, the maximum joint eigenvalue of structural Laplacian follows

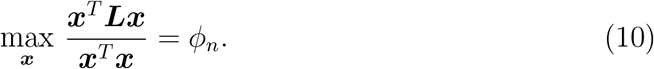

**Proof** : We have ***a***_*i*_, *i* ∈ {1, 2, …, *n*} be joint eigenmodes in *A* such that ***a***_*n*_ is the eigenmode corresponding to the largest joint eigenvalue *n* and ***a***_1_ is the joint eigenmode corresponds to the smallest joint eigenvalue *ϕ*_1_. Suppose *x* = (*c*_1_***a***_1_ + *c*_2_***a***_2_ + … + *n****a***_*n*_), for some constants {*c*_1_, *c*_2_, …, *c*_*n*_}. Then

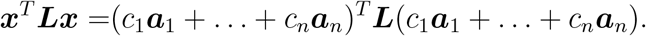

Using the properties of joint eigenmodes ***La***_*i*_ = *ϕ*_*i*_***a***_*i*_ for *i ∈* {1, 2, …, *n*}, we get

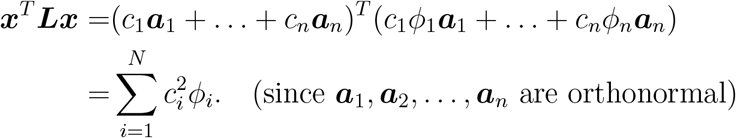

Similarly,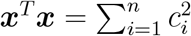. Therefore,

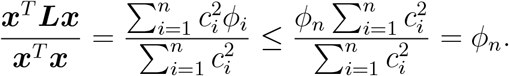

### Lemma 2

Given a symmetric matrix S with non-negative entries, and its diagonal degree matrix *D*, and the matrix 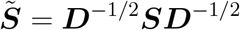, then both 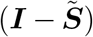 and 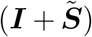 are positive semi-definite (PSD) matrices.

**Proof** : For any given ***x*** ∈ ℝ^*n*^, spanned the joint eigenmodes in *𝒜* we can write:

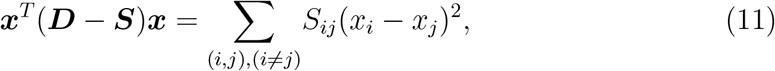

where *S*_*ij*_ = *S*_*ji*_ using the symmetric properties of ***S***. By construction of structural connectivity, each entry of *S* is non-negative real number. Therefore

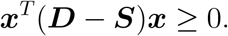

This proves that (***D*** *−****S***) is a positive semi-definite matrix: (***D*** *−****S***) ≽ 0. We use the properties of eigenvalues of multiplication of matrices [61]. Since ***D*** is positive definite matrix, ***D***^*−*1*/*2^ is also positive definite. Therefore

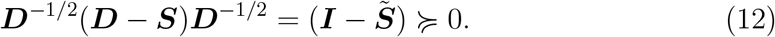

Similar to (11), we have

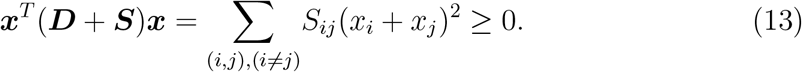

Using the properties of eigenvalues of matrix multiplication,

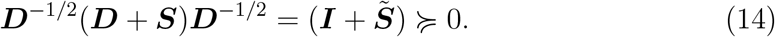

### 6.1. Proof of Theorem 1

The lower bound on eigenvalues of ***L*** directly follows from (12) in Lemma 2. It proves that 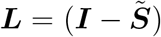 is a PSD matrix. Therefore, the minimum joint eigenvalue of ***L*** follows *ϕ*_1_ *≥* 0.

To show the upper bound of eigenvalues of ***L***, we recall (14) in Lemma 2 that 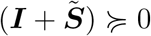. For any *x*, spanned the joint eigenmodes in *𝒜*, we have

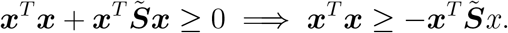

By adding a positive quantity ***x***^*T*^ ***x*** to both sides:

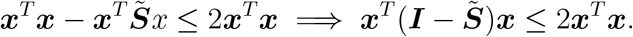

By further simplification,

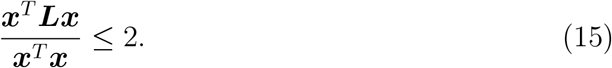

Combining (10) in Lemma 1 and (15), we obtain *ϕ*_*n*_ *≤* 2.

### 6.2. Proof of Theorem 2

We show here that each functional connectome is a positive semi-definite matrix. Without loss of generality, we note that functional connectome [62, 50] basically measures the second moment matrix of a temporal signal (/random variable) ***z*** ∈ ℝ^*n×*1^. In particular,

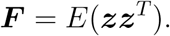

Suppose there are *T* time-point at each ROI is used to estimate the connectivity. Then, 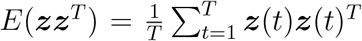. Now, for any random vector ***b*** ∈ ℝ^*n×*1^, we can write

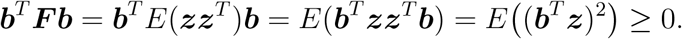

Note that ***b***^*T*^ ***F b*** is the expectation of the square of the scalar random variable 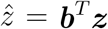. Therefore, for any eigenmodes in 𝒜, the respective eigen spectrum is a nonnegative number.

